# Resistance to aztreonam in combination with non-β-lactam β-lactamase inhibitors due to the layering of mechanisms in *Escherichia coli* identified following mixed culture selection

**DOI:** 10.1101/615336

**Authors:** Ching Hei Phoebe Cheung, Punyawee Dulyayangkul, Kate J. Heesom, Matthew B. Avison

## Abstract

Using mixed culture selection, we show how reduced envelope permeability, reduced target-site affinity, and increased β-lactamase production layer to confer aztreonam/β-lactamase inhibitor resistance in *Escherichia coli.* We report a clinical isolate producing CTX-M-15 and CMY-4, lacking OmpF, and carrying a PBP3 mutation. It is resistant to aztreonam plus the inhibitors avibactam, relebactam and vaborbactam. Mobilisation of *bla*_SHV-12_ into this isolate generated a derivative additionally resistant to aztreonam plus the bicyclic boronate inhibitors 2 and taniborbactam.

## Text

The β-lactam-based class A β-lactamase inhibitors clavulanic acid and tazobactam are widely administered with penicillin derivatives and have had decades of clinical success (1,2). However, class C and D β-lactamases are not affected by these inhibitors, and nor are class B, metallo-β-lactamases, which provide resistance to a wide range of β-lactams including carbapenems (2). Non-β-lactam based β-lactamase inhibitors recently introduced into clinical practice are avibactam, relebactam and vaborbactam, and these have a wider spectrum of activity than clavulanic acid or tazobactam, including many class C and some class D enzymes, but they do not inhibit class B enzymes (2). Brem *et al.* showed that bicyclic boronates are potent inhibitors of both serine-β-lactamases (classes A, C and D) and the most common subclass of metallo-β-lactamases, subclass B1 (3). Another bicyclic boronate “cross-class” β-lactamase inhibitor, VNRX-5133 is in phase 3 clinical trials and known as taniborbactam (4).

Metallo-β-lactamase producers are usually susceptible to aztreonam, because monobactams are very poor substrates for these enzymes (1). However, aztreonam is broken down by class A ESBLs and class C β-lactamases, so isolates carrying a metallo-β-lactamase and one of these serine enzymes are typically aztreonam resistant. Therefore, a combination of aztreonam with a serine-β-lactamase inhibitor has been proposed to kill bacteria carrying a metallo-β-lactamase and a serine enzyme that hydrolyses aztreonam. The combination currently receiving most attention, and undergoing clinical trials is aztreonam/avibactam (3).

A study of 328 *Escherichia coli* resistant to third generation cephalosporins showed an aztreonam/avibactam MIC of ≤ 0.25/4 μg.ml^-1^ in all except one isolate that carried CTX-M-15 (class A ESBL) and hyper-produced the chromosomal AmpC (class C enzyme) against which the aztreonam/avibactam MIC was 1/4 μg.ml^-1^ (5). Following on from this observation, we tested for reduced susceptibility (based on the CLSI broth microdilution protocol [6]) to aztreonam in combination with avibactam (4 μg.ml^-1^) and other recently released serine β-lactamase inhibitors relebactam (4 μg.ml^-1^) and vaborbactam (8 μg.ml^-1^) against *E. coli* producing a CTX-M and a plasmid mediated AmpC (pAmpC) enzyme. In our collection of sequenced human *E. coli,* we located two relevant isolates. Urinary isolate 9969 (ST69) (7) carries genes encoding CTX-M-15 and CMY-60 (pAmpC). Bloodstream isolate N16 (ST101) (a gift from Prof Tim Walsh, University of Oxford) carries genes encoding CTX-M-15 and CMY-4. Against isolate 9969, all three inhibitors (purchased from MedChem Express) brought the aztreonam MIC back into the susceptible range, based on CSLI breakpoints (8). In contrast N16 was resistant to aztreonam in the presence of all three inhibitors (**Table 1**).

**Table 1.**
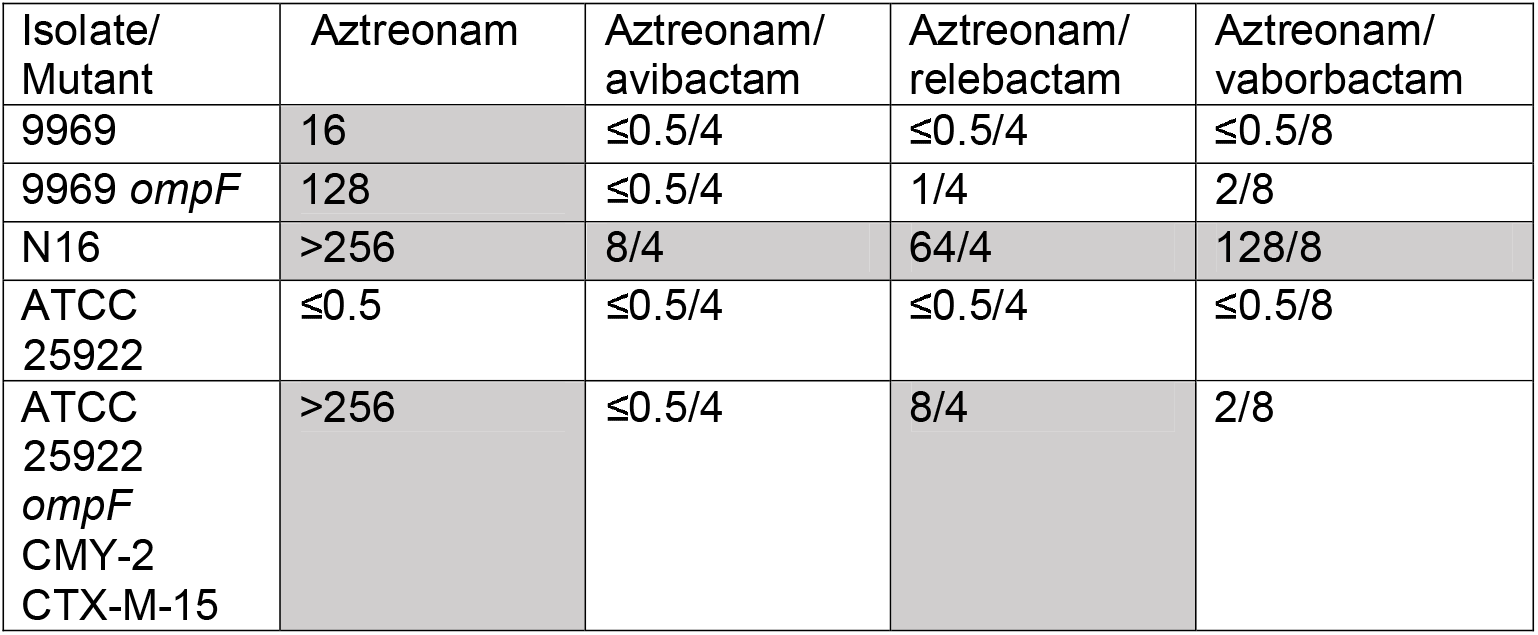
MIC (μg.ml^-1^) of Aztreonam with and without serine β-lactamase inhibitors against *E. coli* isolates and mutant derivatives.

| Isolate/<br>Mutant | Aztreonam | Aztreonam/<br>avibactam | Aztreonam/<br>relebactam | Aztreonam/<br>vaborbactam |
| --- | --- | --- | --- | --- |
| 9969 | 16 | $\leq 0.5/4$ | $\leq 0.5/4$ | $\leq 0.5/8$ |
| 9969 *ompF* | 128 | $\leq 0.5/4$ | 1/4 | 2/8 |
| N16 | >256 | 8/4 | 64/4 | 128/8 |
| ATCC<br>25922 | $\leq 0.5$ | $\leq 0.5/4$ | $\leq 0.5/4$ | $\leq 0.5/8$ |
| ATCC<br>25922<br>*ompF*<br>CMY-2<br>CTX-M-15 | >256 | $\leq 0.5/4$ | 8/4 | 2/8 |

Analysis of whole genome sequence (WGS) data revealed an 8 bp insertion, leading to a frameshift in the *ompF* porin gene in N16. In 9969, *ompF* is wild-type, so we hypothesised that this loss of OmpF is the reason why N16 is resistant to aztreonam plus the inhibitors. To test this, we insertionally inactivated *ompF* in 9969 using pKNOCK suicide gene replacement (9). An *ompF* DNA fragment was amplified with Phusion High-Fidelity DNA Polymerase (NEB, UK) from *E. coli* 9969 genomic DNA using primers EC-*ompF*-KO-FW (5’ CAA<u>GGATCC</u>TGATGGCCTGAACTTC 3’) with BamHI restriction site, underlined, and EC-*ompF*-KO-RV (5’ CAA<u>GTCGAC</u>TTCAGACCAGTAGCC 3’) with SalI restriction site, underlined, digested with BamHI and SalI and ligated into pKNOCK-GM at the same sites generating pKNOCK-GM::*ompF.* This was transferred into 9969 by conjugation from *E. coli* BW20767. Mutants were selected using cefoxitin (5 μg.mL^-1^) and gentamicin (5 μg.mL^-1^) and confirmed by PCR using primers *EC-ompF-F* (5’ ATGATGAAGCGCAATAAT 3’) and BT543 (5’ TGACGCGTCCTCGGTAC 3’). Whilst disruption of *ompF* in isolate 9969 did increase the MIC of aztreonam, and of aztreonam with either relebactam or vaborbactam, it did not detectably raise the MIC of aztreonam in the presence of avibactam, nor did it confer resistance to aztreonam in the presence of any inhibitor (**Table 1**). We hypothesised that this susceptibility difference between N16 (an *ompF* mutant) and 9969*ompF* was due to higher level production of CMY and/or CTX-M in N16 versus 9969, and proteomics analysis, performed as described by us previously (10,11), showed slightly higher enzyme production in N16 versus 9969 (**Table 2**). To test whether producing high levels of CMY/CTX-M in an *ompF* mutant confers aztreonam/inhibitor resistance, susceptible *E. coli* ATCC25922 was transformed with the multi-copy plasmids pYT(*bla*_CTX-M-15_) and pSU18(*bla*_CMY-2_), made as previously described (10,12). We the generated an *ompF* mutant derivative of this ATCC25922 transformant as above, with the mutant selected on gentamicin (5 μg.mL^-1^), cefoxitin (5 μg.mL^-1^) and kanamycin (20 μg.mL^-1^). Proteomics confirmed the average abundance of CMY was 0.48 per unit of ribosomal proteins for N16 and 3.44 for ATCC25922*ompF*(CTX-M-15,CMY-2) (n=3 for both). For CTX-M-15, the average abundance was 0.44 per unit of ribosomal proteins for N16 and 1.27 for ATCC25922*ompF*(CTX-M-15,CMY-2), which was resistant to aztreonam/relebactam but not aztreonam/vaborbactam; the MIC of aztreonam in the presence of avibactam against ATCC25922*ompF*(CTX-M-15,CMY-2) remained very low (**Table 1**). Therefore, we conclude that slightly higher levels of CMY/CTX-M production is not the reason for aztreonam/avibactam resistance in isolate N16, even when combined with OmpF porin loss and there must be an additional mechanism. Further WGS analysis revealed a 12 nt tandem duplication in *ftsI* in N16, predicted to cause the insertion of the amino acid sequence YRIN after proline 333 in PBP3. This aztreonam target site mutation has previously been seen in clinical isolates, and associated with elevated MICs of aztreonam, with and without avibactam (13–15). Accordingly, the collective effects of CTX-M-15 and CMY production, loss of OmpF and a target site mutation in PBP3 explain why N16 is resistant to aztreonam in the presence of all three serine β-lactamase inhibitors tested.

**Table 2.**
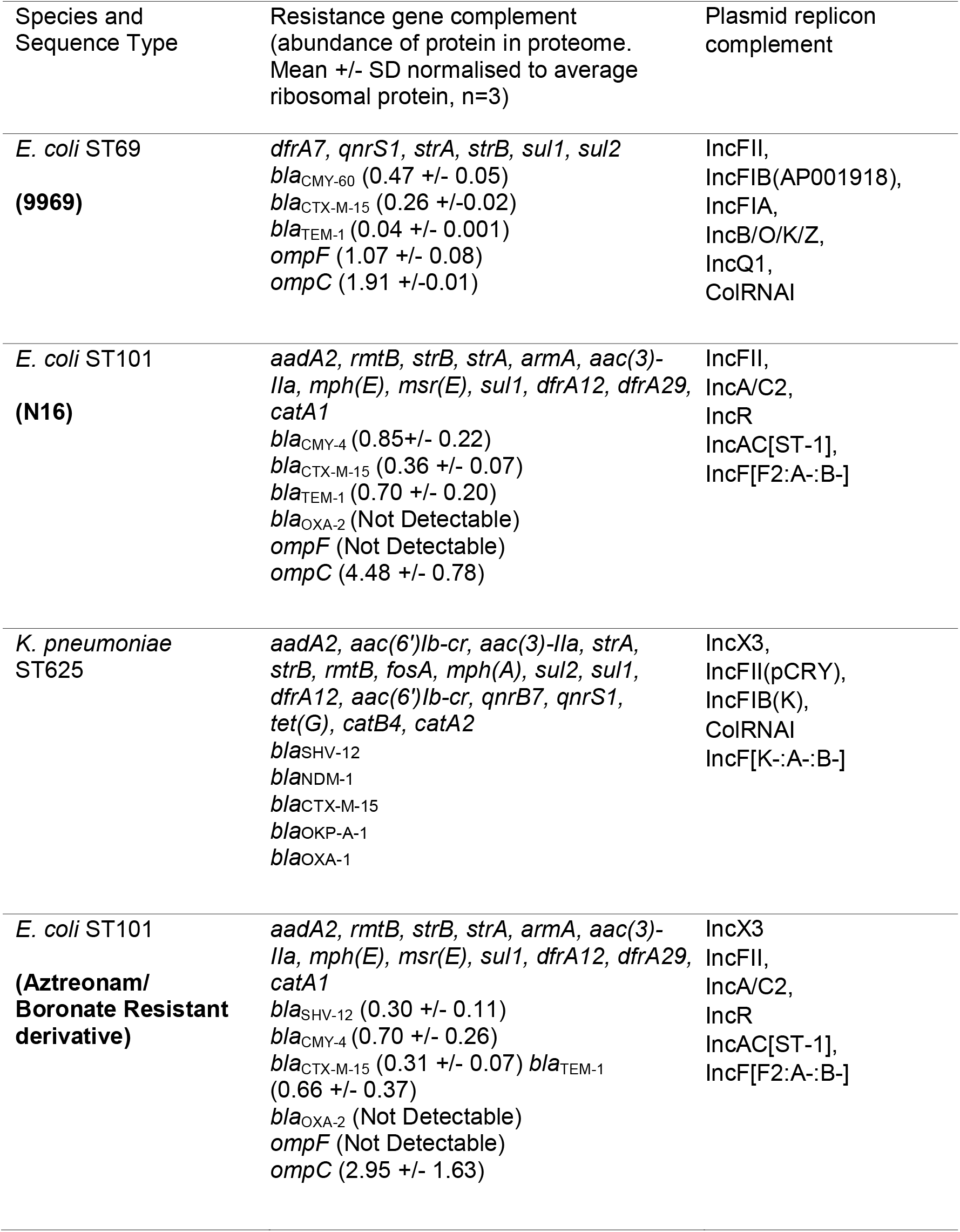
Genotypic and Phenotypic Properties of *E. coli* and *K. pneumoniae* isolates.

The cross-class bicyclic boronate inhibitor **2** inhibits pAmpCs and CTX-M β-lactamases (3,16). The aztreonam/avibactam resistant *E. coli* isolate N16 was susceptible (MIC = 4 μg.ml^-1^) to aztreonam in the presence of 10 μg.ml^-1^ of inhibitor **2**, synthesized according to the literature protocol (3) and kindly provided by Prof. Chris Schofield, University of Oxford. We then spent some considerable time trying to select N16 point mutants resistant to aztreonam/**2** but failed. During these attempts, N16 and a *K. pneumoniae* ST265 bloodstream isolate (also a gift from Prof Tim Walsh) were mistakenly inoculated into the same bottle containing cation adjusted Muller-Hinton broth (CA-MHB) and grown overnight without any antibiotic selection. **Table 2** lists the resistance gene and plasmid replicon carriage status of the two isolates. One hundred microlitres of the mixed overnight culture was plated onto Muller Hinton agar containing aztreonam at increasing concentrations plus inhibitor **2** at a fixed concentration of 10 μg.ml^-1^. Profuse growth was seen on all plates up to 16 μg.ml^-1^ aztreonam, which is defined as resistant by CLSI (8). One aztreonam/**2** resistant *E. coli* colony was selected as representative; the MIC of aztreonam in the presence of 4 μg.ml^-1^ of another cross-class bicyclic boronate inhibitor, taniborbactam (5) was 32 μg.ml^-1^, confirming that this N16 derivative is more generally aztreonam/inhibitor resistant.

Dye accumulation assays (17,18) showed that *E. coli* N16 and its resistant derivative had similar envelope permeability (**Figure 1**) and proteomic analysis showed no significant difference in the abundances of key porin and efflux pump proteins (**Table 2**). WGS revealed that the complement of β-lactamases and plasmid replicon types in the resistant N16 derivative had increased compared with N16; an SHV-12 encoding IncX3 plasmid had moved from the *K. pneumoniae* isolate into N16 during co-culture (**Table 2**). SHV-12 was produced at high levels in the N16 resistant derivative, and the abundances of the other resident β-lactamases was not significantly different from N16 (**Table 2**). Notably, whilst IncX3 plasmids have previously been seen to carry *bla*_SHV-12_ and *bla*_NDM_ (19), the *bla*_NDM_ gene located in our *K. pneumoniae* donor isolate did not co-transfer with *bla*_SHV-12_ into N16 (**Table 2**) and it has been reported previously that *bla*_SHV-12_ has been identified on IncX3 plasmids lacking *bla*_NDM_ in *E. coli* (20).

**Figure 1:**
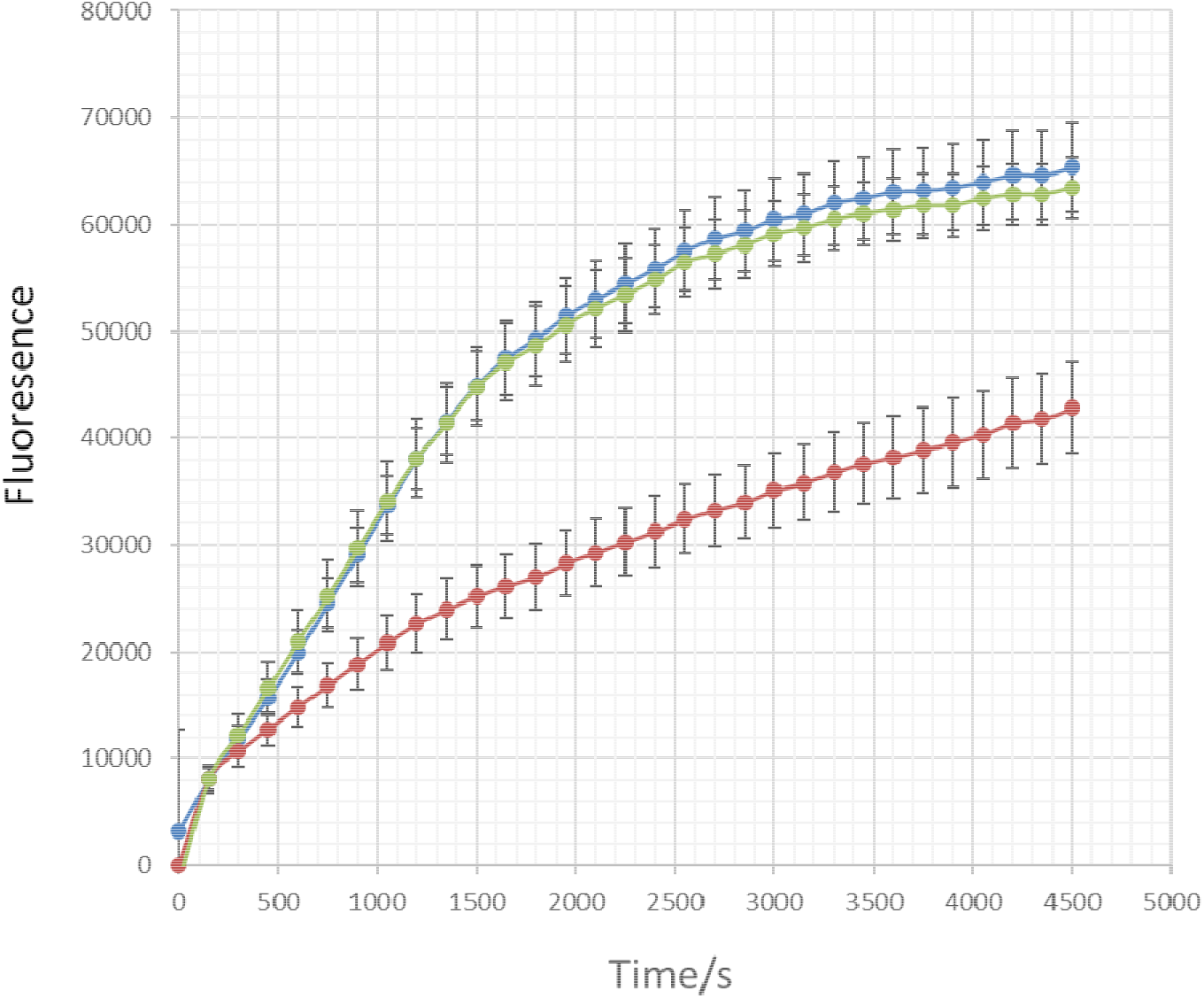
The accumulation of H33342 dye over a 30 cycle (4500 s) incubation period by *K. pneumoniae* and *E. coli* isolates. Red line, *K. pneumoniae* ST625 donor isolate; blue line, *E. coli* N16; green line, *E. coli* N16 resistant derivative. In each case, fluorescence of cells incubated with the dye is presented as an absolute value after each cycle. Each line shows mean data for three biological replicates with 8 technical replicates in each. Error bars define the standard error of the mean.

Whilst *E. coli* N16 (and its resistant derivative) carry genes for *bla*_TEM-1_ and *bla*_OXA_-_2_, only the former was detectably expressed (**Table 2**). WGS confirmed that *bla*_OXA_._2_ is chromosomally located (so is single copy), the third gene cassette in the integron (so is distant from the integron common promoter) and the integron promoter is of the weakest known designation (21), explaining its low-level expression.

We conclude, therefore, that the in the absence of OmpF and with an aztreonam target site mutation in PBP3, the production of three β-lactamases (CMY-2, CTX-M-15, and SHV-12) that all hydrolyse aztreonam (1) from enzyme classes that bind bicyclic boronates (3,22) perhaps with a contribution from a resident TEM-1, which also binds bicyclic boronates (3) collectively overcome utility of aztreonam/**2** and aztreonam/taniborbactam.

The fortuitous finding reported here has significant implications for the future of research into β-lactam/β-lactamase inhibitor resistance. It is known that β-lactamase hyperproduction - following gene duplication, promoter mutation, or mutations that stabilise the enzyme - can overcome β-lactam/β-lactamase inhibitor combinations (e.g., amoxicillin/clavulanate or ceftazidime/avibactam) (1,2,23) but clearly another way of increasing the abundance of β-lactamase activity in a cell is to acquire an additional β-lactamase gene from a neighbouring bacterium with similar substrate profile, as we have found here. This could never be seen when testing individual isolates for their ability to generate resistant derivatives; either in the lab or using in vivo infection models. However, in the real world, mixed populations of bacteria are found, increasing the potential for resistance to coalesce in one member of the population via horizontal gene transfer from the “β-lactamase-ome” of the population. Two final findings are relevant to note. First, the bicyclic boronate taniborbactam has been partnered with cefepime, not aztreonam in clinical development (5). The MIC of cefepime against *E. coli* isolate N16 in the presence of taniborbactam was 4/4 μg.ml^-1^, and this did not rise upon acquisition of *bla*_SHV-12_ in the aztreonam/taniborbactam resistant N16 derivative. This MIC is in CLSI’s new “susceptible-dose dependent” range for cefepime (8). Second, prior to clinical approval, aztreonam/avibactam as a combination is created in the clinical setting by dosing aztreonam alongside ceftazidime/avibactam. Using checkerboard assays (**Fig. 2**) we noted that the aztreonam/avibactam resistant N16 and its aztreonam/taniborbactam resistant derivative are both susceptible (though sitting at the breakpoint) to ceftazidime/avibactam (MIC = 8/4 μg.ml^-1^) even without aztreonam. The MIC versus N16 falls slightly to 4/4 μg.ml^-1^ in the presence of a breakpoint concentration of aztreonam (4 μg.ml^-1^) but not against the aztreonam/taniborbactam resistant N16 derivative. Accordingly, in addition to the use of carbapenems (provided a metallo-β-lactamase is not also present), it may be possible to treat infections with isolates having the same combination of β-lactam resistance mechanisms as N16 or its aztreonam/bicyclic boronate resistant derivative by using ceftazidime/avibactam (with or without aztreonam) or cefepime/taniborbactam but correct dosing may be difficult to achieve because the observed MICs are so close to the breakpoints.

**Figure 2:**
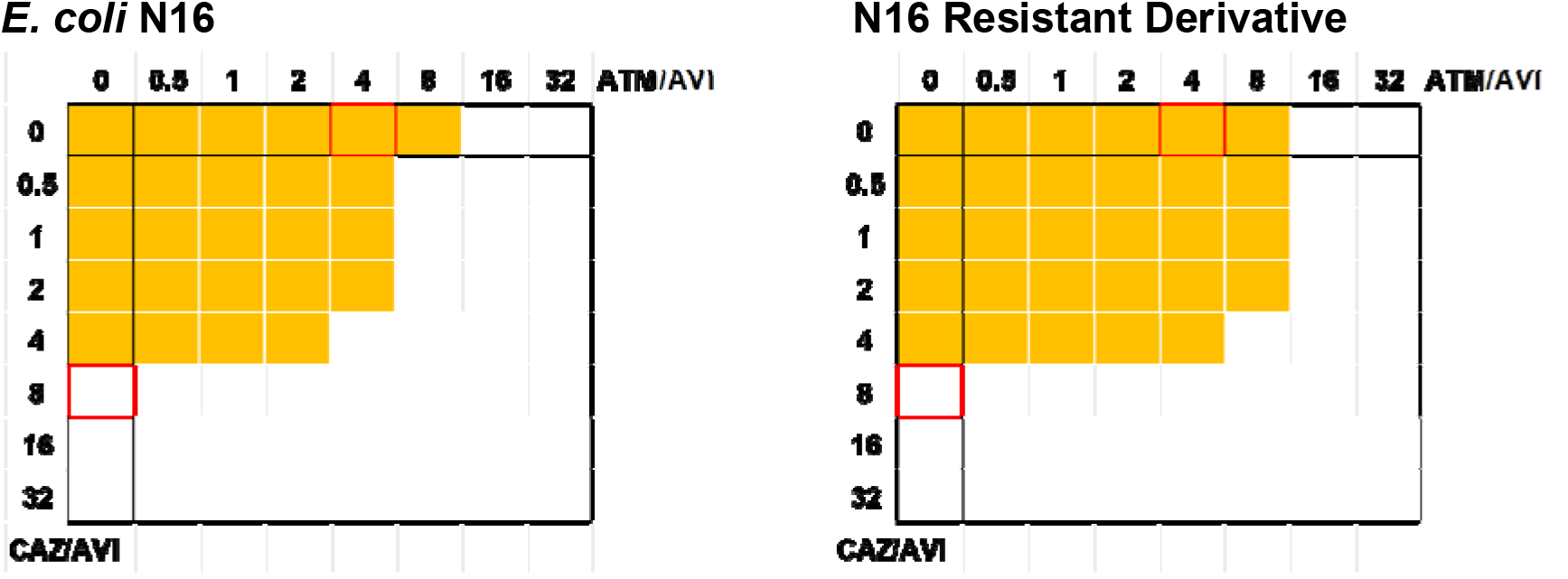
Checkerboard assays for ceftazidime aztreonam in the presence of avibactam. Each image represents triplicate assays for an 8×8 array of wells in a 96-well plate. All wells contained CA-MHB including avibactam (4 μg.mL^-1^). A serial dilution of aztreonam (ATM, x-axis) and ceftazidime (CAZ, y-axis) was created from 32 μg.mL^-1^ in each plate as recorded. All wells were inoculated with a suspension of bacteria, made as per CLSI microtiter MIC guidelines (6), and the plate was incubated at 37°C for 20 h. Growth was recorded by measuring OD_600_ and growth above background (broth) is recorded as a yellow block. Growth at 8 μg.mL^-1^ ceftazidime and 4 μg.mL^-1^ aztreonam (this position indicated in red) in the presence of avibactam defines resistance based on CLSI breakpoints (8). Bacterial suspensions used were N16, left hand image, and the N16 derivative selected for aztreonam/bicyclic boronate resistance, right hand image.

## Acknowledgments

This work was funded by grants MR/N013646/1 to M.B.A. and K.J.H. and MR/S004769/1 to M.B.A. from the Antimicrobial Resistance Cross Council Initiative supported by the seven United Kingdom research councils and the National Institute for Health Research. Additional funding came via a bequest from the estate of the late Professor Graham Ayliffe. Genome sequencing was provided by MicrobesNG (http://www.microbesng.uk/).

**We declare no conflicts of interest.**

